# Incorporating uniparental markers and demographic information in kinship analysis

**DOI:** 10.1101/2022.05.05.490843

**Authors:** Jin-Xian Liu, Meng-Yu Li

## Abstract

Knowledge of kinship relations between members of wild populations is of great importance in ecological and conservation genetic studies. The bi-parentally inherited autosomal markers has been the Golden Standard in kinship analysis. However, analysis of kin relationship can be challenging in wild populations. The uni-parentally inherited markers and population demographic information can be helpful for identifying false-positive in kinship analysis. Here we showed how incorporating uniparental genetic and demographic information can improve the correct classification rate of kinship analyses by reanalyzing data of a recent study published in Science Advances. The application of next generation high-throughput sequencing to address fundamental ecological questions is of immense benefit to the field of molecular ecology, which could also generate uniparentally inherited organelle genomes together with nuclear data. We strongly recommended that uniparental genetic markers and demographic information be seriously considered in kinship analyses of wild populations.

## MAIN TEXT

Knowledge of kinship relations between members of wild populations is of great importance in ecological and conservation genetic studies with potentially far-reaching implications (Weir, Anderson, & Hepler, 2006; Blouin, 2003; Flanagan & Jones, 2019; Speed, & Balding, 2015). Kinship analyses can provide indispensable information for the study of dispersal (Jasper et al., 2019; Schunter et al., 2014), mating systems (Bentzen et al., 2001), variance of reproductive success among individuals (Christie et al. 2010; Liu & Ely 2009), inbreeding avoidance (Hargrove et al., 2021), kin recognition, and kin selection (O’Corry-Crowe et al., 2020) as well as aiding the management of endangered populations (Escoda, Fernández-González & Castresana, 2019; Kanno, Vokoun & Letcher, 2011). However, analysis of kin relationship can be challenging in wild populations (Weir, Anderson & Hepler, 2006). Kinship can be estimated on the basis of genetic similarity among individuals of a population, reflecting the proportion of homologous alleles shared between 2 individuals due to identity-by-descent from a common ancestor (Blouin, 2003). Estimates of genetic relatedness may be inaccurate because they require the assumption that founders are outbred and unrelated. High levels of inbreeding may affect relatedness and therefore kinship inference (Wang, 2014). Even for individuals whose parents are not inbred, account needs to be taken of ‘background relatedness’ that is due to evolutionary history in a population (Weir, Anderson & Hepler, 2006). So the assessment of kinship among members of wild animal populations is difficult in the absence of detailed multigenerational pedigrees and background relatedness (Stadele & Vigilant, 2016). Parent-offspring relationships can be determined with higher confidence than other relationships by comparing the genetic information of an offspring and supposed parents under the simple Mendel’s laws (Flanagan & Jones, 2019). However, it becomes markedly more challenging in situations where neither parent is known by field observation. In species for which the offspring of unsampled parents can be expected to contain groups of full and/or half siblings, sibship reconstruction is a powerful tool for identifying related individuals (Jones et al. 2010). However, sibship reconstruction for populations with complex kinship structures, which present in species with promiscuous mating systems, long life spans, and overlapping generations, can also lead to high error rates (Stadele & Vigilant, 2016). In addition, genetic data could suffer from a high rate of genotyping errors such as allelic dropouts and false alleles in situations such as data from low-coverage next-generation sequencing, noninvasive samples, and highly degraded samples (Blouin 2003; Wang et al. 2019). Use of such data without accounting for mistyping properly could also lead to inaccurate or incorrect inferences of kin relationships.

The bi-parentally inherited autosomal markers, e.g. SNPs or microsatellites, has been the Golden Standard in kinship analysis because of their high discriminatory power in kinship inference (Goudet et al., 2018; Weinman, Solomon & Rubenstein, 2015). The uni-parentally inherited (sex-specific inherited) markers are usually applied to cases where autosomal genotypes cannot be obtained due to technical constraints. The disadvantage of uniparental markers is that their profiles are usually not individual specific, resulting in a much lower discrimination power (Kayser, 2007). The uniparental markers yields limited information compared with autosomal markers and does not allow one to obtain a complete reconstruction of possible kin relationships (Vai et al., 2020). However, the inheritance patterns of the uniparentally inherited markers make them powerful additions to kinship analyses using bi-parentally inherited autosomal data by clearly identifying false-positive assignments of kinship with Mendelian incompatibilities (Kopps et al., 2015). For example, individuals not sharing their maternally inherited haplotypes could not be assigned to the category full sibs even if they had high likelihood. In addition to genetic data, demographic information can also be helpful for identifying false-positive. For example, individuals of different age class could not be assigned to the category full sibs.

High-throughput next-generation sequencing platforms can generate high-density genome-wide data in a relatively short time and at an affordable price. With rapidly increasing applications of genome-wide SNPs in model and non-model species, dense markers has been used in kinship analyses of wild populations by taking advantage of the high informative power (Escoda, Fernández-González & Castresana, 2019; Jasper et al., 2019; Martin, Lipps & Gibbs, 2021). Next-generation sequencing projects can also generate organelle DNA in addition to nuclear DNA, which makes the sequence analysis of entire organelle genomes feasible for kinship analyses (Al-Nakeeb et al. 2017). Although the use of high-density, genome-wide molecular markers can enable reasonably accurate assignment of individuals to high-order groupings (Goudet et al., 2018; (Phillips, García-Magariños, Salas, Carracedo & Lareu, 2012), the organelle DNA sequences generated together by next-generation sequencing platforms could be expected to provide a nice chance to excluded some false-positive results. To illustrate how incorporating demographic and uniparental genetic information can improve the correct classification rate of kinship analyses, here we reanalyzed the data of a recent study (Vendramli et al. 2021) published in Science Advances by incorporating the maternally inherited mitochondrial DNA and age information.

Based on sibship reconstruction and simulation analyses, Vendramli et al. (2021) reported an extreme reproductive sweepstake event together with collective larval dispersal in the highly abundant shallow water Antarctic limpet, *Nacella concinna,*with which they explained the chaotic genetic patchiness observed in the Rose Garden 1999. Vendramli et al. (2021) draw their conclusions under a complete open population frame, which implies that the recruitment of local populations of *N. concinna* were exclusively from other sources. Based on the results of sibship reconstruction, their forward genetic simulations implicated the Rose Garden 1999 age class most likely originated from an extreme sweepstake reproductive success of a single female residing within one stack in an unknown source population. Then at least 95% of the offspring from the extreme reproductive sweepstake in the source population dispersed collectively to Rose Garden. The findings were quite intriguing, however, both the extreme sweepstake reproductive success and strong collective dispersal are only theoretically sound from their forward simulations. For organisms with great fecundity and high mortality in early life stages, the need to match reproductive activity with environmental conditions conducive to spawning, fertilization, larval development and recruitment may result in extreme variance in reproductive success among individuals (Hedgecock & Pudovkin, 2011). Inherent to the sweepstake reproductive success hypothesis is the idea that variance in reproductive success among breeders is extremely high. However, spawning of *N. concinna* is highly synchronous among all breeding adults in a given population and occurs in a very narrow timeframe no longer than 24 hours (Picken & Allan, 1983; Stanwell-Smith & Clarke, 1998). The larvae produced by the breeding group of a specific population during such a short period of time may form a single cohort and meet similar environmental conditions, which apparently does not facilitate the occurrence of extreme variance in reproductive success among individuals. Furthermore, larvae transported far from natal populations by ocean currents usually suffer high mortality along the way (Becker, Levin, Fodrie, & McMillan, 2007; Thorson, 1950). So successful recruitment of most larvae to Rose Garden from the extreme reproductive sweepstake in an unknown source population via collective dispersal with low mortality could be logistically challenging.

If the Rose Garden 1999 sample were originated from an extreme sweepstake reproductive success of a single female as suggested, the haplotype diversity of maternally inherited mitochondrial sequences for the Rose Garden 1999 sample should be extremely lower than those in other populations. More specifically, only one mitochondrial haplotype shared by the offspring of the single super female should be expected in Rose Garden 1999. The Restriction site-associated DNA sequencing (RAD-seq) conducted in the original study (Vendrami et al., 2021) was performed on both nuclear and mitochondrial genome simultaneously. There are two Eco RI recognition sites in the complete, 16761 base pairs long, mitochondrial genome of *N. concinna* (GenBank accession: KT990126.1), at nucleotide positions from 10645 to 10650 and from 14984 to 14989 respectively. However, authors of the original study ignored the mitochondrial sequences in their data, which could validate their conclusion of the extreme sweepstake reproductive success directly.

Here we tested a specific prediction of the extreme sweepstake reproductive success proposed in Vendramli et al. (2021), sharing of a single mitochondrial haplotype by the offspring of the single reproductive sucessful female.The clean paired-end reads of each individual were mapped to the reference mitochondrial genome. After removing contaminated and low-quality individual dataset, consensus haploid mitochondrial sequence was called for 132 individuals from 14 samples. We got two fragments of mitochondrial sequences with a total length of 1645 base pairs corresponding to the two Eco RI recognition sites. One fragment corresponds to 10235-11066 in the reference mitochondrial genome including partial ND1, tRNA-Leu and partial16S ribosomal RNA. The other fragment corresponds to 14568-15380 of the reference mitochondrial genome including partial COX3, tRNA-Arg, tRNA-Asn, and partial ND3. In addition, a third Eco RI recognition site originating from a C to T transition at 6384 of the mitochondrial genome changing GAACTC to GAATTC was also detected in six individuals. The average depth of coverage per nucleotide position of the 1645 bp per individual ranged from 42 X to 443 X and were lager than 100 X for most (121/91%) individuals, consistent with the high copy number of mitochondrial genome per cell (Table S1). Comparison of the 132 mitochondrial sequences revealed 20 distinct haplotypes defined by 22 polymorphic sites with 19 transitions, and 3 single-base deletions (Table S2). Most of the haplotypes (12/60%) were singletons represented by only a single individual. The remaining eight haplotypes were shared by multiple populations (Table S3). The median-joining network of the 20 haplotypes observed revealed a shallow star-like topology with a dominant haplotype H1 (68 individuals, ~53%) in the center (Fig. 1), suggesting a shallow phylogeographic history, which was consistent with previous study with a partial fragment of the COI gene in the western Antarctic Peninsula (González-Wevar, David, & Poulin, 2011). The number of haplotypes observed per population ranged from 3 in four samples (EB15, RG99, Do 99, and SI99) to 7 in RG15. The haplotype diversity *(h)* per population ranged from 0.38 in EB15 to 0.93 in RG15 (Table 1). Low levels of nucleotide diversity were detected in all localities.

**Figure 1.**
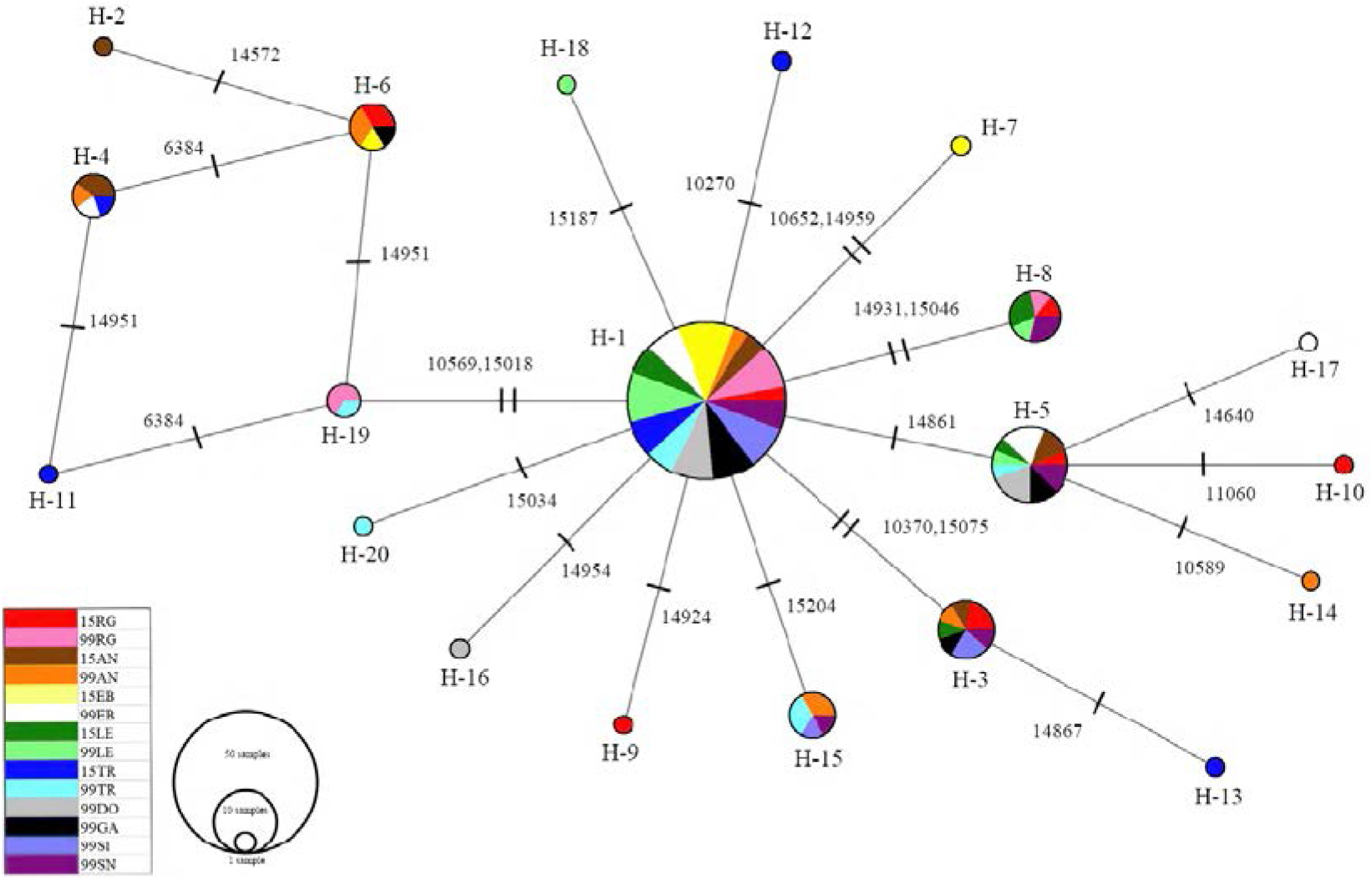
Median-joining network for 20 mt haplotypes of *Nacella concinna.* Haplotypes are represented by circles, the sizes of which are proportional to the number of individuals. Different colors represent geographic distribution. Mutational steps between haplotypes are indicated by hatch marks

**Table 1.**
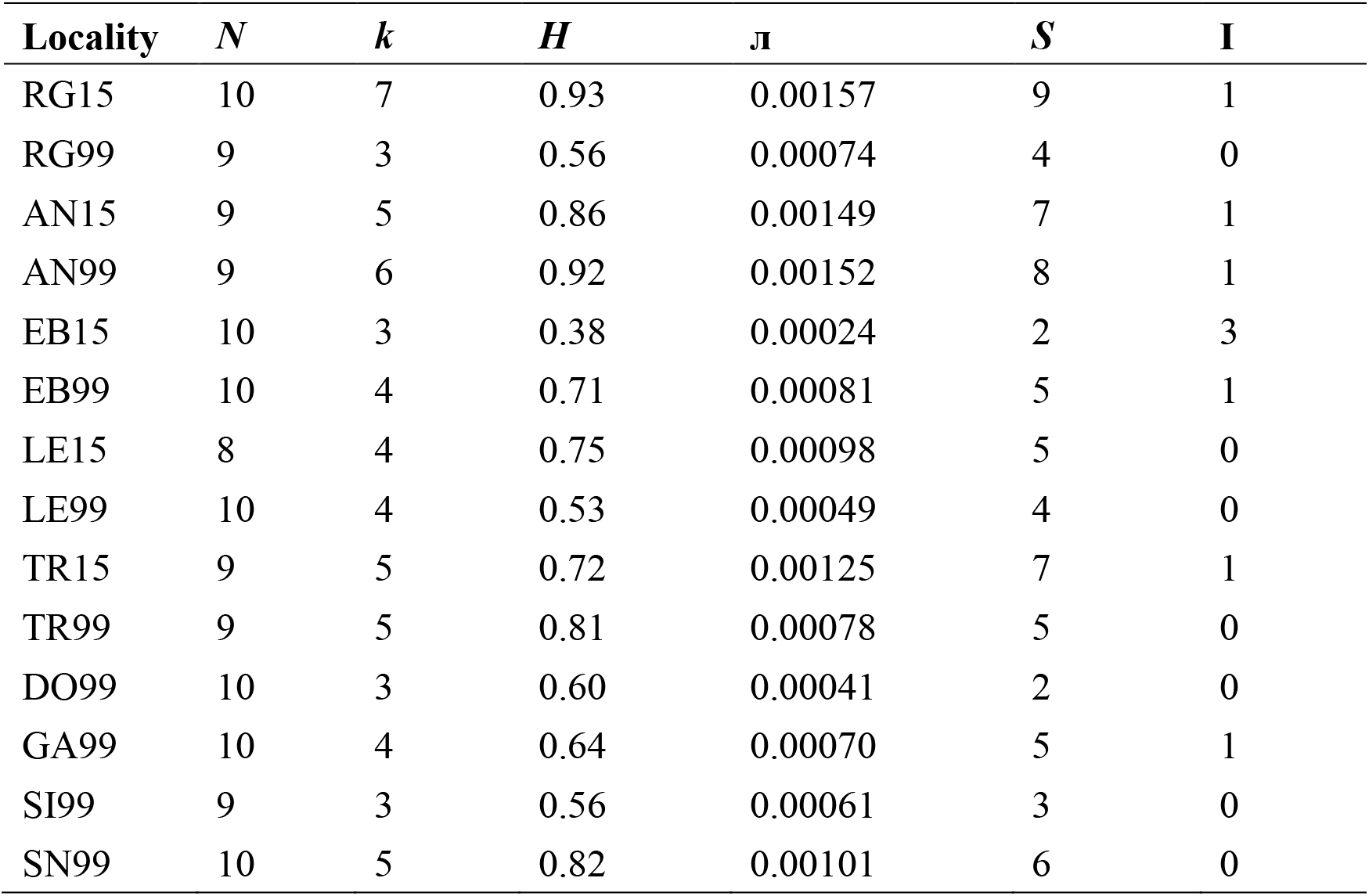
Summary of genetic diversity indices for samples of *Nacella concinna. N*: number of sampled specimens; *k:* number of haplotypes detected; *H:* haplotype diversity; Л: nucleotide diversity; *S:* polymorphic sites; I: number of indels.

In contrast to the extreme sweepstake reproductive success claimed in the original study (Vendrami et al., 2021) in the Rose Garden 1999, three mitochondrial haplotypes were detected in only nine individuals of the Rose Garden 1999 sample by using the 1646 base pairs of mitochondrial sequences. Six of the nine individuals were found with the most dominant haplotype H1 that shared by 68 individuals among all the populations. Two individuals were found with a subdominant haplotype H19 shared by three individuals from Ryder Bay. The last individual was detected with another subdominant haplotype H8 shared by seven individuals from five individuals. If each of the three mitochondrial haplotypes detected corresponded to one spawning female, then at least three females had contributed to the recruitment of the Rose Garden 1999 cohort. However, considering that all the three haplotypes were dominant or subdominant ones, it is highly possible that the individuals shared the same mitochondrial haplotype in the Rose Garden 1999 sample could be offspring of different spawning females with the same dominant or subdominant haplotype, as evidenced in other samples with unrelated individuals but shared haplotypes. Furthermore, the three haplotypes were detected with only nine individuals of the highly abundant *N. concinna* and with partial mitochondrial sequences of 1646 base pairs, which is only about one tenth of the whole mitochondrial genome. If more individuals and more sequences of the mitochondrial genome (e.g. whole mitochondrial genome) were analyzed, it could be of high probability that more mitochondrial haplotypes could be found in the Rose Garden 1999 cohort. Indeed, previous research with a partial fragment (663 bp) of the mitochondrial COI gene, which was not targeted by the RAD-Seq in Vendramli et al. (Vendrami et al., 2021), for populations of *N. concinna* in the western Antarctic Peninsula revealed 16 haplotypes among 160 individuals, indicating informative polymorphisms in other parts of the mitochondrial genome and possibly higher haplotype diversity considering more mitochondrial sequences (González-Wevar et al., 2011).

Clearly, by analyzing the maternally inherited mitochondrial sequences in the original RAD-Seq data of Vendramli et al. (2021), the number and distribution of mitochondrial haplotypes in the Rose Garden 1999 cohort indicated that multiple spawning females, at least three and highly possible many more, had contributed to the recruitment the Rose Garden 1999 age class. Clearly, the extreme sweepstake reproductive success claimed by Vendramli et al. (2021) for the Rose Garden 1999 cohort was not supported by the maternally inherited mitochondrial sequences in their own dataset, let alone the collective dispersal of larvae from the extreme sweepstake reproductive success in a source population.

The important evidence supporting an extreme sweepstake event in Vendramli et al. (2021) is that Rose Garden in 1999 was exclusively represented by full-and half-siblings, which was assessed based on the relatedness coefficients. Any relatedness between individuals occurs against a background level of relatedness in the population, either as a consequence of inbreeding or by belonging to the same population (Weir, Anderson, & Hepler, 2006). However, some relatedness measures for use with molecular data assume that the individuals themselves are not inbred, as the approach (Manichaikul et al., 2010) that adopted by Vendramli et al. (2021).

Given that the mitochondrial data did not support the sweepstake hypothesis, the close kinships assessed among individuals in the Rose Garden 1999 cohort could possibly reflect a background relatedness resulting from high degree of inbreeding, the mating of individuals closely related by ancestry. Vendramli et al. (2021) explained the chaotic genetic patchiness in the Rose Garden 1999 cohort with an extreme sweepstake reproductive success that supported by a suite of correlated genomic signatures including locally reduced genetic diversity, substantially elevated genomic relatedness, and locally elevated linkage disequilibrium. However, inbreeding can also be expected to lead to low polymorphism, extensive linkage disequilibrium, low effective population size, and high population subdivision (Charlesworth, 2003). Indeed, the number of mitochondrial haplotypes, haplotype diversity, and nucleotide diversity were lower in the Rose Garden 1999 sample than in the Rose Garden 2015 sample, suggesting signals of possible inbreeding.

Furthermore, a critical prerequisite for the extreme sweepstake reproductive success that proposed by Vendramli et al. (2021) is that all the limpets analyzed in Rose Garden 1999 were from the same age cohort. To ensure that, Vendramli et al. restricted their sampling to limpets with shells between 20 and 30 mm, which they thought corresponded to animals approximately 10 years of age. However, typical of polar marine invertebrates, growth rate for *N. concinna* is slow, which makes it difficult to follow the growth of individual cohorts or year classes (Clarke, Prothero-Thomas, Beaumont, Chapman, & Brey, 2004; Picken, 1980). A previous study found that the most abundant size group of *Nacella concinna* comprised individuals with sizes from 20 to 30 mm, and the von Bertalanffy growth curve demonstrated that individuals with shell size from 20 to 30 mm included multiple age classes (Brêthes, Ferreyra, & de la Vega, 1994). Furthermore, the annual shell growth derived from mark and recapture techniques was slow, with annual increment less than 3 mm for most individuals with shell size from 20 to 30 mm (Clarke et al., 2004). Based on all these evidences, it was highly possible that multiple age cohorts might exist among individuals of *N. concinna* with shell size between 20 and 30 mm. So the so-called full-and half-siblings observed in Rose Garden 1999 by Vendramli et al. (2021) might only reflect close relatedness among individuals, but not real full- and half-siblings as suggested.

By incorporating demographic and uniparental genetic information, our results did not support sweepstake reproductive success and collective dispersal in the Rose Garden 1999 sample of *N. concinna.* Similarly, cohesive dispersal over extensive periods (4–6 month) was also suggested in the splitnose rockfish *(Sebastes diploproa)* by genetic relatedness analysis with nuclear microsatellites, which indicated that 11.6% of the recruits were siblings in a single recruitment pulse (Ottmann et al., 2016). However, further sequencing of the mitochondrial control region demonstrated that the juvenile samples consisted two different rockfish species, which inflated estimates of relatedness within dyads containing individuals from the same species (Ottmann et al., 2017). Thus, the original analysis with microsatellite data did not have evidence to support the long-term cohesive dispersal as claimed.

In summary, although bi-parental genetic markers play the major role in kinship analysis, incorporating demographic and uniparental genetic information can improve the correct classification rate of kinship analyses by reducing false positives. The application of next generation high-throughput sequencing to address fundamental ecological questions is of immense benefit to the field of molecular ecology. Although theoretically, even distant kin relationships can be accurately classified when a large number of markers, linkage information, or whole-genome sequence data can be attained (Kling et al., 2012; Li et al., 2014), our results suggested that misclassification can still happen due to complex background of wild populations. Since the sequences of uniparental organelle DNA could be routinely generated together with nuclear DNA by the next generation high-throughput sequencing, we strongly recommended that these uniparental genetic information be seriously considered in kinship analyses of wild populations. In particular, because of its high abundance in cells, the assembly of organelle genomes can be obtained even with only low-coverage next generation sequencing data (Rasheed et al., 2017), which could be of particular importance for kinship studies with low-coverage NGS data, where genotyping error and misclassification rate could be high (Wang et al. 2019). In addition, knowledge of population demographic information such as age and sex are also very important in kinship analyses, even in the genomic era.

## Data availability

Data are available in the original paper (3), The BAM files used for consensus sequences extraction and the aligned FASTA file of the sequences analyzed in this study are available at figshare:https://10.6084/m9.figshare.16937386.

## Author contributions

J-X.L. supervised the study. M-Y.L. and J-X.L.analyzed the data, J-X.L. wrote the manuscript with contribution of M-Y.L.

## Supplementary materials

This manuscript has supplementary materials with details of Materials and Methods and supplementary tables.

