## Supplementary text and tables for "Incorporating uniparental markers and demographic information in kinship analysis"

### This PDF file includes:

Materials and methods

Tables S1 to S4

References

**Materials and Methods**

**Mitochondrial Sequence Extraction**

The quality-filtered RAD sequencing data of Vendramli et al. (2021) were downloaded from the Sequence Read Archive (SRA accession number: PRJNA728555, <https://ncbi.nlm.nih.gov/sra/?term=PRJNA728555>). The mitochondrial genome of *N concinna* (GenBank accession: KT990126.1) was used as reference for mapping. Sequencing reads were de-multiplexed and mapped against the reference mitochondrial genome using BWA v.0.1.17 (Li, 2013) with the mem algorithm and default settings. The resulting BAM files were then indexed, cleaned and sorted using SAMtools version 0.1.19 (Li et al., 2009). Extraction of the mapped sequence was performed using mpileup algorithm in SAMtools with the following settings: --ff 0*400 --ff 0*100 --rf 0*2 -C50 -q 20 and vcfutils.pl in BCFtools version 0.1.19 (Li et al., 2009) with minimum depth set as 3.

The extraction of mitochondrial sequences failed in three individuals (SRR14478765, SRR14478791, SRR14478855). In total, 137 mitochondiral sequences were aligned with ClustalW and manually edited using BIOEDIT version 7.0.5.3 (Hall, 1999). After removing five low quality and individuals with contamination, we got the haploid consensus sequences of two fragments in 132 individuals corresponding to the two Eco RI recognition sites in the reference mitochondrial genome. After trimming, a total of 1645 base pairs of the two Eco RI recognition sites associated fragments were retained for all the 132 individuals. In addition, a third Eco RI recognition site was also detected in six individuals. Only the variable nucleotide position in the third Eco RI recognition site were used for subsequent analysis. In total, 1646 base pairs of combined mitochondrial sequences were used for downstream analysis. The depth of coverage per nucleotide position in the two Eco RI recognition site associated fragments in all individuals were obtained from BAM files using htsbox pileup algorithm (<https://github.com/lh3/htsbox>) with minimum depth set as 3.

**Molecular Diversity and Haplotype Network**

The mitochondrial sequences were converted into Arlequin format using DnaSP v.5 (Librado & Rozas, 2009). The number of polymorphic sites (*S*), haplotype diversity (*H*) and nucleotide diversity (**л**) were calculated with DnaSP v.5. Median-joining haplotype network was constructed with program Network, version 4.6 (http://www.fluxus-engineering.com/) (Bandelt, Forster, & Rohl, 1999).

### Table S1.

Summary of statistics for sequencing and mapping of 132 individuals of *Nacella concinna*.

| **Sample name** | **Geo_loc_name** | **Pop** | **Average**  **depth** | **Bases** |
| --- | --- | --- | --- | --- |
| SRR14478745 | Antarctica: Ryder Bay - Anchorage Island - Rose Garden | RG15 | 201 | 1.19 G |
| SRR14478746 | Antarctica: Ryder Bay - Trolval | TR15 | 131 | 880.14 M |
| SRR14478747 | Antarctica: Ryder Bay - Anchorage Island - Rose Garden | RG15 | 187 | 1.27 G |
| SRR14478748 | Antarctica: Ryder Bay - Anchorage Island - Rose Garden | RG15 | 164 | 756.80 M |
| SRR14478749 | Antarctica: Ryder Bay - Anchorage Island - Rose Garden | RG15 | 197 | 1.26 G |
| SRR14478750 | Antarctica: Ryder Bay - Anchorage Island - Rose Garden | RG15 | 175 | 908.39 M |
| SRR14478751 | Antarctica: Ryder Bay - Anchorage Island - Rose Garden | RG15 | 321 | 1.42 G |
| SRR14478752 | Antarctica: Ryder Bay - Anchorage Island - Rose Garden | RG15 | 210 | 1.20 G |
| SRR14478753 | Antarctica: Ryder Bay - Anchorage Island - Rose Garden | RG15 | 236 | 1.27 G |
| SRR14478754 | Antarctica: Ryder Bay - Anchorage Island - Rose Garden | RG15 | 100 | 618.66 M |
| SRR14478755 | Antarctica: Ryder Bay - Anchorage Island - Rose Garden | RG15 | 252 | 1.22 G |
| SRR14478756 | Antarctica: Ryder Bay - Anchorage Island North | AN99 | 112 | 1.12 G |
| SRR14478757 | Antarctica: Ryder Bay - Leonie Island | LE15 | 284 | 1.00 G |
| SRR14478758 | Antarctica: Ryder Bay - Leonie Island | LE15 | 358 | 2.22 G |
| SRR14478759 | Antarctica: Ryder Bay - Leonie Island | LE15 | 333 | 1.66 G |
| SRR14478760 | Antarctica: Ryder Bay - Leonie Island | LE15 | 423 | 1.94 G |
| SRR14478761 | Antarctica: Ryder Bay - Leonie Island | LE15 | 443 | 2.37 G |
| SRR14478763 | Antarctica: Ryder Bay - Leonie Island | LE15 | 243 | 1.19 G |
| SRR14478764 | Antarctica: Ryder Bay - Leonie Island | LE15 | 168 | 1.06 G |
| SRR14478766 | Antarctica: Ryder Bay - Leonie Island | LE15 | 151 | 1.04 G |
| SRR14478768 | Antarctica: Ryder Bay - East Beach | EB15 | 361 | 1.23 G |
| SRR14478769 | Antarctica: Ryder Bay - East Beach | EB15 | 230 | 1.37 G |
| SRR14478770 | Antarctica: Ryder Bay - East Beach | EB15 | 87 | 553.26 M |
| SRR14478771 | Antarctica: Ryder Bay - East Beach | EB15 | 277 | 1.27 G |
| SRR14478772 | Antarctica: Ryder Bay - East Beach | EB15 | 202 | 1.24 G |
| SRR14478773 | Antarctica: Ryder Bay - East Beach | EB15 | 133 | 733.69 M |
| SRR14478774 | Antarctica: Ryder Bay - Trolval | TR15 | 265 | 1.67 G |
| SRR14478775 | Antarctica: Dobrowolski Island | DO99 | 225 | 1.08 G |
| SRR14478776 | Antarctica: Ryder Bay - Anchorage Island North | AN99 | 151 | 1.20 G |
| SRR14478777 | Antarctica: Ryder Bay - Anchorage Island North | AN99 | 243 | 1.47 G |
| SRR14478778 | Antarctica: Ryder Bay - East Beach | EB15 | 137 | 700.23 M |
| SRR14478779 | Antarctica: Ryder Bay - East Beach | EB15 | 198 | 1.22 G |
| SRR14478780 | Antarctica: Ryder Bay - East Beach | EB15 | 252 | 1.49 G |
| SRR14478781 | Antarctica: Ryder Bay - East Beach | EB15 | 178 | 865.03 M |
| SRR14478782 | Antarctica: Ryder Bay - Anchorage Island North | AN99 | 127 | 1.27 G |
| SRR14478783 | Antarctica: Ryder Bay - Anchorage Island North | AN15 | 129 | 782.62 M |
| SRR14478784 | Antarctica: Ryder Bay - Anchorage Island North | AN15 | 104 | 709.59 M |
| SRR14478785 | Antarctica: Ryder Bay - Anchorage Island North | AN15 | 90 | 663.65 M |

| SRR14478786 | Antarctica: Ryder Bay - Anchorage Island North | AN15 | 112 | 794.37 M |
| --- | --- | --- | --- | --- |
| SRR14478787 | Antarctica: Ryder Bay - Anchorage Island North | AN15 | 130 | 888.12 M |
| SRR14478788 | Antarctica: Ryder Bay - Anchorage Island North | AN15 | 137 | 765.44 M |
| SRR14478789 | Antarctica: Ryder Bay - Anchorage Island North | AN15 | 165 | 1.02 G |
| SRR14478790 | Antarctica: Ryder Bay - Anchorage Island North | AN15 | 179 | 898.43 M |
| SRR14478792 | Antarctica: Ryder Bay - Anchorage Island North | AN15 | 84 | 697.65 M |
| SRR14478793 | Antarctica: Ryder Bay - Anchorage Island North | AN99 | 108 | 1.13 G |
| SRR14478794 | Antarctica: Snow Island | SN99 | 278 | 1.78 G |
| SRR14478795 | Antarctica: Snow Island | SN99 | 199 | 1.12 G |
| SRR14478796 | Antarctica: Snow Island | SN99 | 112 | 818.57 M |
| SRR14478797 | Antarctica: Snow Island | SN99 | 240 | 1.27 G |
| SRR14478798 | Antarctica: Snow Island | SN99 | 317 | 1.47 G |
| SRR14478799 | Antarctica: Snow Island | SN99 | 155 | 1.27 G |
| SRR14478800 | Antarctica: Snow Island | SN99 | 284 | 1.48 G |
| SRR14478801 | Antarctica: Snow Island | SN99 | 226 | 1.29 G |
| SRR14478802 | Antarctica: Snow Island | SN99 | 142 | 1.14 G |
| SRR14478803 | Antarctica: Snow Island | SN99 | 180 | 1.18 G |
| SRR14478804 | Antarctica: Ryder Bay - Anchorage Island North | AN99 | 139 | 1.41 G |
| SRR14478805 | Antarctica: Ryder Bay - Leonie Island | LE99 | 197 | 1.25 G |
| SRR14478806 | Antarctica: Ryder Bay - Leonie Island | LE99 | 210 | 1.32 G |
| SRR14478807 | Antarctica: Ryder Bay - Leonie Island | LE99 | 269 | 1.25 G |
| SRR14478808 | Antarctica: Ryder Bay - Leonie Island | LE99 | 257 | 1.54 G |
| SRR14478809 | Antarctica: Ryder Bay - Leonie Island | LE99 | 182 | 1.12 G |
| SRR14478810 | Antarctica: Ryder Bay - Leonie Island | LE99 | 171 | 1.18 G |
| SRR14478811 | Antarctica: Ryder Bay - Leonie Island | LE99 | 146 | 1.12 G |
| SRR14478812 | Antarctica: Ryder Bay - Leonie Island | LE99 | 183 | 1.36 G |
| SRR14478813 | Antarctica: Ryder Bay - Leonie Island | LE99 | 213 | 1.05 G |
| SRR14478814 | Antarctica: Ryder Bay - Leonie Island | LE99 | 242 | 1.57 G |
| SRR14478815 | Antarctica: Ryder Bay - Anchorage Island North | AN99 | 204 | 1.34 G |
| SRR14478816 | Antarctica: Galindez Island | GA99 | 203 | 1.55 G |
| SRR14478817 | Antarctica: Galindez Island | GA99 | 240 | 1.38 G |
| SRR14478818 | Antarctica: Galindez Island | GA99 | 105 | 927.04 M |
| SRR14478819 | Antarctica: Galindez Island | GA99 | 223 | 1.47 G |
| SRR14478820 | Antarctica: Galindez Island | GA99 | 142 | 1.17 G |
| SRR14478821 | Antarctica: Galindez Island | GA99 | 281 | 1.79 G |
| SRR14478822 | Antarctica: Galindez Island | GA99 | 281 | 1.15 G |
| SRR14478823 | Antarctica: Galindez Island | GA99 | 226 | 1.45 G |
| SRR14478824 | Antarctica: Galindez Island | GA99 | 235 | 1.31 G |
| SRR14478825 | Antarctica: Galindez Island | GA99 | 42 | 478.44 M |
| SRR14478826 | Antarctica: Ryder Bay - Anchorage Island North | AN99 | 148 | 1.41 G |
| SRR14478827 | Antarctica: Ryder Bay - East Beach | EB99 | 82 | 1.46 G |
| SRR14478828 | Antarctica: Ryder Bay - East Beach | EB99 | 125 | 1.35 G |
| SRR14478829 | Antarctica: Ryder Bay - East Beach | EB99 | 104 | 1.10 G |
| SRR14478830 | Antarctica: Ryder Bay - East Beach | EB99 | 85 | 1.39 G |

| SRR14478831 | Antarctica: Ryder Bay - East Beach | EB99 | 128 | 1.60 G |
| --- | --- | --- | --- | --- |
| SRR14478832 | Antarctica: Ryder Bay - East Beach | EB99 | 154 | 1.92 G |
| SRR14478833 | Antarctica: Ryder Bay - East Beach | EB99 | 143 | 1.44 G |
| SRR14478834 | Antarctica: Ryder Bay - East Beach | EB99 | 89 | 1.23 G |
| SRR14478835 | Antarctica: Ryder Bay - East Beach | EB99 | 154 | 1.63 G |
| SRR14478836 | Antarctica: Ryder Bay - East Beach | EB99 | 130 | 1.41 G |
| SRR14478837 | Antarctica: Ryder Bay - Anchorage Island North | AN99 | 146 | 1.01 G |
| SRR14478838 | Antarctica: Dobrowolski Island | DO99 | 216 | 1.16 G |
| SRR14478839 | Antarctica: Dobrowolski Island | DO99 | 287 | 1.42 G |
| SRR14478840 | Antarctica: Dobrowolski Island | DO99 | 260 | 1.39 G |
| SRR14478841 | Antarctica: Dobrowolski Island | DO99 | 192 | 1.24 G |
| SRR14478842 | Antarctica: Dobrowolski Island | DO99 | 169 | 1.11 G |
| SRR14478843 | Antarctica: Dobrowolski Island | DO99 | 211 | 1.44 G |
| SRR14478844 | Antarctica: Ryder Bay - Trolval | TR99 | 251 | 1.61 G |
| SRR14478846 | Antarctica: Ryder Bay - Trolval | TR99 | 149 | 1.16 G |
| SRR14478847 | Antarctica: Ryder Bay - Trolval | TR99 | 198 | 1.28 G |
| SRR14478848 | Antarctica: Ryder Bay - Trolval | TR99 | 165 | 1.15 G |
| SRR14478849 | Antarctica: Ryder Bay - Trolval | TR99 | 259 | 1.30 G |
| SRR14478850 | Antarctica: Ryder Bay - Trolval | TR99 | 96 | 806.68 M |
| SRR14478851 | Antarctica: Ryder Bay - Trolval | TR99 | 122 | 1.02 G |
| SRR14478852 | Antarctica: Ryder Bay - Trolval | TR99 | 196 | 1.42 G |
| SRR14478853 | Antarctica: Ryder Bay - Trolval | TR99 | 174 | 941.07 M |
| SRR14478854 | Antarctica: Dobrowolski Island | DO99 | 350 | 2.57 G |
| SRR14478856 | Antarctica: Signy Island | SI99 | 142 | 965.85 M |
| SRR14478857 | Antarctica: Signy Island | SI99 | 282 | 1.23 G |
| SRR14478858 | Antarctica: Signy Island | SI99 | 147 | 1.31 G |
| SRR14478859 | Antarctica: Signy Island | SI99 | 216 | 1.67 G |
| SRR14478860 | Antarctica: Signy Island | SI99 | 181 | 1.24 G |
| SRR14478861 | Antarctica: Signy Island | SI99 | 169 | 969.60 M |
| SRR14478862 | Antarctica: Signy Island | SI99 | 164 | 1.04 G |
| SRR14478863 | Antarctica: Signy Island | SI99 | 96 | 885.52 M |
| SRR14478864 | Antarctica: Signy Island | SI99 | 246 | 1.49 G |
| SRR14478865 | Antarctica: Dobrowolski Island | DO99 | 252 | 2.54 G |
| SRR14478866 | Antarctica: Ryder Bay - Anchorage Island - Rose Garden | RG99 | 165 | 1.05 G |
| SRR14478867 | Antarctica: Ryder Bay - Anchorage Island - Rose Garden | RG99 | 267 | 1.50 G |
| SRR14478868 | Antarctica: Ryder Bay - Anchorage Island - Rose Garden | RG99 | 209 | 1.28 G |
| SRR14478870 | Antarctica: Ryder Bay - Anchorage Island - Rose Garden | RG99 | 182 | 1.06 G |
| SRR14478871 | Antarctica: Ryder Bay - Anchorage Island - Rose Garden | RG99 | 177 | 1.06 G |
| SRR14478872 | Antarctica: Ryder Bay - Anchorage Island - Rose Garden | RG99 | 232 | 1.26 G |
| SRR14478873 | Antarctica: Ryder Bay - Anchorage Island - Rose Garden | RG99 | 218 | 1.19 G |
| SRR14478874 | Antarctica: Ryder Bay - Anchorage Island - Rose Garden | RG99 | 275 | 1.43 G |
| SRR14478875 | Antarctica: Ryder Bay - Anchorage Island - Rose Garden | RG99 | 236 | 970.76 M |
| SRR14478876 | Antarctica: Dobrowolski Island | DO99 | 222 | 1.53 G |
| SRR14478877 | Antarctica: Ryder Bay - Trolval | TR15 | 414 | 1.71 G |

| SRR14478879 | Antarctica: Ryder Bay - Trolval | TR15 | 137 | 979.21 M |
| --- | --- | --- | --- | --- |
| SRR14478880 | Antarctica: Ryder Bay - Trolval | TR15 | 146 | 962.71 M |
| SRR14478881 | Antarctica: Ryder Bay - Trolval | TR15 | 47 | 240.13 M |
| SRR14478882 | Antarctica: Ryder Bay - Trolval | TR15 | 104 | 680.27 M |
| SRR14478883 | Antarctica: Ryder Bay - Trolval | TR15 | 134 | 583.18 M |
| SRR14478884 | Antarctica: Ryder Bay - Trolval | TR15 | 119 | 814.37 M |

### Table S2.

Variable sites defining 20 distinct haplotypes among 132 individuals from 14 populaitons of *Nacella concinna*. ID in ref: position of the variable sites in the reference mitochondrial genome.

| **ID in ref** | 6384 | 10270 | 10370 | 10569 | 10589 | 10652 | 11060 | 14572 | 14640 | 14861 | 14867 | 14924 | 14931 | 14951 | 14954 | 14959 | 15018 | 15034 | 15046 | 15075 | 15187 | 15204 |
| --- | --- | --- | --- | --- | --- | --- | --- | --- | --- | --- | --- | --- | --- | --- | --- | --- | --- | --- | --- | --- | --- | --- |
| H_1 | C | T | C | T | T | T | A | A | T | A | T | A | T | T | T | T | T | G | A | T | A | C |
| H_2 | . | . | . | C | . | . | . | G | . | . | . | . | . | - | . | . | C | . | . | . | . | . |
| H_3 | . | . | T | . | . | . | . | . | . | . | . | . | . | . | . | . | . | . | . | C | . | . |
| H_4 | T | . | . | C | . | . | . | . | . | . | . | . | . | - | . | . | C | . | . | . | . | . |
| H_5 | . | . | . | . | . | . | . | . | . | G | . | . | . | . | . | . | . | . | . | . | . | . |
| H_6 | . | . | . | C | . | . | . | . | . | . | . | . | . | - | . | . | C | . | . | . | . | . |
| H_7 | . | . | . | . | . | - | . | . | . | . | . | . | . | . | . | - | . | . | . | . | . | . |
| H_8 | . | . | . | . | . | . | . | . | . | . | . | . | C | . | . | . | . | . | G | . | . | . |
| H_9 | . | . | . | . | . | . | . | . | . | . | . | G | . | . | . | . | . | . | . | . | . | . |
| H_10 | . | . | . | . | . | . | G | . | . | G | . | . | . | . | . | . | . | . | . | . | . | . |
| H_11 | T | . | . | C | . | . | . | . | . | . | . | . | . | . | . | . | C | . | . | . | . | . |
| H_12 | . | C | . | . | . | . | . | . | . | . | . | . | . | . | . | . | . | . | . | . | . | . |
| H_13 | . | . | T | . | . | . | . | . | . | . | C | . | . | . | . | . | . | . | . | C | . | . |
| H_14 | . | . | . | . | C | . | . | . | . | G | . | . | . | . | . | . | . | . | . | . | . | . |
| H_15 | . | . | . | . | . | . | . | . | . | . | . | . | . | . | . | . | . | . | . | . | . | T |
| H_16 | . | . | . | . | . | . | . | . | . | . | . | . | . | . | C | . | . | . | . | . | . | . |
| H_17 | . | . | . | . | . | . | . | . | C | G | . | . | . | . | . | . | . | . | . | . | . | . |
| H_18 | . | . | . | . | . | . | . | . | . | . | . | . | . | . | . | . | . | . | . | . | G | . |
| H_19 | . | . | . | C | . | . | . | . | . | . | . | . | . | . | . | . | C | . | . | . | . | . |
| H_20 | . | . | . | . | . | . | . | . | . | . | . | . | . | . | . | . | . | A | . | . | . | . |

### Table S3.

Distribution of the 20 mitochondrial haplotypes among 14 *Nacella concinna* populations.

| **POP** | **H_1** | **H_2** | **H_3** | **H_4** | **H_5** | **H_6** | **H_7** | **H_8** | **H_9** | **H_10** | **H_11** | **H_12** | **H_13** | **H_14** | **H_15** | **H_16** | **H_17** | **H_18** | **H_19** | **H_20** |
| --- | --- | --- | --- | --- | --- | --- | --- | --- | --- | --- | --- | --- | --- | --- | --- | --- | --- | --- | --- | --- |
| **RG15** | 2 | 0 | 2 | 0 | 1 | 2 | 0 | 1 | 1 | 1 | 0 | 0 | 0 | 0 | 0 | 0 | 0 | 0 | 0 | 0 |
| **RG99** | 6 | 0 | 0 | 0 | 0 | 0 | 0 | 1 | 0 | 0 | 0 | 0 | 0 | 0 | 0 | 0 | 0 | 0 | 2 | 0 |
| **AN15** | 3 | 1 | 1 | 2 | 2 | 0 | 0 | 0 | 0 | 0 | 0 | 0 | 0 | 0 | 0 | 0 | 0 | 0 | 0 | 0 |
| **AN99** | 2 | 0 | 1 | 1 | 0 | 2 | 0 | 0 | 0 | 0 | 0 | 0 | 0 | 1 | 2 | 0 | 0 | 0 | 0 | 0 |
| **EB15** | 8 | 0 | 0 | 0 | 0 | 1 | 1 | 0 | 0 | 0 | 0 | 0 | 0 | 0 | 0 | 0 | 0 | 0 | 0 | 0 |
| **EB99** | 5 | 0 | 0 | 1 | 3 | 0 | 0 | 0 | 0 | 0 | 0 | 0 | 0 | 0 | 0 | 0 | 1 | 0 | 0 | 0 |
| **LE15** | 4 | 0 | 1 | 0 | 1 | 0 | 0 | 2 | 0 | 0 | 0 | 0 | 0 | 0 | 0 | 0 | 0 | 0 | 0 | 0 |
| **LE99** | 7 | 0 | 0 | 0 | 1 | 0 | 0 | 1 | 0 | 0 | 0 | 0 | 0 | 0 | 0 | 0 | 0 | 1 | 0 | 0 |
| **TR15** | 5 | 0 | 0 | 1 | 0 | 0 | 0 | 0 | 0 | 0 | 1 | 1 | 1 | 0 | 0 | 0 | 0 | 0 | 0 | 0 |
| **TR99** | 4 | 0 | 0 | 0 | 1 | 0 | 0 | 0 | 0 | 0 | 0 | 0 | 0 | 0 | 2 | 0 | 0 | 0 | 1 | 1 |
| **DO99** | 6 | 0 | 0 | 0 | 3 | 0 | 0 | 0 | 0 | 0 | 0 | 0 | 0 | 0 | 0 | 1 | 0 | 0 | 0 | 0 |
| **GA99** | 6 | 0 | 1 | 0 | 2 | 1 | 0 | 0 | 0 | 0 | 0 | 0 | 0 | 0 | 0 | 0 | 0 | 0 | 0 | 0 |
| **SI99** | 6 | 0 | 2 | 0 | 0 | 0 | 0 | 0 | 0 | 0 | 0 | 0 | 0 | 0 | 1 | 0 | 0 | 0 | 0 | 0 |
| **SN99** | 4 | 0 | 1 | 0 | 2 | 0 | 0 | 2 | 0 | 0 | 0 | 0 | 0 | 0 | 1 | 0 | 0 | 0 | 0 | 0 |
| **Total** | 68 | 1 | 9 | 5 | 16 | 6 | 1 | 7 | 1 | 1 | 1 | 1 | 1 | 1 | 6 | 1 | 1 | 1 | 3 | 1 |
